# Earlier finish of motor planning in the premotor cortex predicts faster motor command in the primary motor cortex: human intracranial EEG evidence

**DOI:** 10.1101/2024.03.05.583627

**Authors:** Jing Xia, Deshan Gong, Biao Han, Qiang Guo, Gereon R. Fink, Silvia Daun, Qi Chen

## Abstract

The human motor system has a hierarchical control during finger movements. The non-primary motor cortex (premotor cortex, PM, and supplementary motor area, SMA) organizes motor planning, while the primary motor cortex (M1) is responsible for motor execution. We utilized the human intracranial EEG’s high temporal and spatial resolution to investigate how the temporal dynamics of the high-gamma neural oscillations in the hierarchically organized motor sub-regions, during both pre-movement planning and motor execution, correlated with reaction times (RTs) in a cued finger movement task. Our results showed that the high-gamma power of PM, SMA, and M1 activated sequentially. More importantly, the sustained high-gamma power activation in the non-primary motor cortex and the peak latency of high-gamma power in M1 had a significant predictive relationship with the RTs. In particular, the faster the activation of the non-primary motor cortex returned to baseline, the faster the motor command in M1, and accordingly the shorter the RTs. Further, pairwise phase coherence between the motor areas showed that the more sustained the connection between the motor areas, the longer the RTs would be. The current findings illustrate the relationship between the temporal profiles of high-gamma power in human motor areas and response performance.

## Introduction

Goal-directed voluntary movements result from sophisticated coordination between motor planning in the higher-order motor cortices and motor command in the primary motor cortex (Rao et al. 1993; Christensen et al. 2007; Afshar et al. 2011; Lara et al. 2018). The primary motor cortex (M1, Brodmann area 4), located at the posterior bank of the precentral gyrus, is a critical cortical structure underlying movement execution (Crone et al. 1998). Specifically, the event-related synchronization (ERS) of high-gamma band (60-140Hz, HGB) rhythms in M1, contralateral to the moving body effectors (Miller et al. 2007), has been shown to underlie motor command (Crone *et al*. 1998; Pfurtscheller et al. 2003; Szurhaj et al. 2005; Fonken et al. 2016). The premotor cortex (PM, lateral Brodmann area 6) and the supplementary motor area (SMA, medial Brodmann area 6) have been associated with preparatory activity before motor command (Churchland, Santhanam, et al. 2006; Churchland, Yu, et al. 2006; Churchland and Shenoy 2007; Nachev et al. 2008; Guo et al. 2014; Li et al. 2015; Hasegawa et al. 2017). Specifically, HGB power in the PM has been linked to tongue and hand movement preparation (Miller *et al*. 2007). Notably, HGB synchronization in the non-primary motor regions occurs during the preparation of contralateral and ipsilateral hand movements (Kermadi et al. 2000; Meyer-Lindenberg et al. 2002; Gonzalez et al. 2006; Wilson et al. 2010). Moreover, the firing rate of PM during motor preparation has been shown to predict subsequent movement speed in monkeys: trials with higher firing rate variability and distribution in the preparation tended to result in longer reaction times (Bastian et al., 2003; Churchland et al., 2006).

However, both anatomical structures and functions of the motor cortex differ notably between humans and monkeys (Schieber 1999; Schieber and Santello 2004). The human motor cortex exhibits a greater diversity of neurons, with different neurons manifesting distinct patterns in controlling finger movements (Kleinschmidt et al. 1997; Schieber 1999; Beisteiner et al. 2001; Kim 2001). The human motor cortex is better suited for highly individualized and fine-grained control of finger movements, whereas the monkey motor cortex is more oriented towards coordinating the movements of multiple finger muscles (Georgopoulos et al. 1999; Schieber and Santello 2004; Ben Hamed et al. 2007). For example, the anatomical organization of M1, featuring both diverging and converging corticospinal output pathways, establishes a neural foundation for complex finger movements. In human M1, a somatotopic gradient is observed, with the thumb representation being more pronounced laterally and the little finger representation more medially. This somatotopic gradient, however, is less apparent in macaque M1 (Schieber 1999; Schieber and Santello 2004). Moreover, distinct patterns of representation activity during finger movements, particularly flexion and extension, have been observed in high-field 7T fMRI experiments conducted on humans compared to spiking recordings in monkeys (Arbuckle et al. 2020).

Given the structural and functional differences in the motor cortex between humans and monkeys, it remains poorly understood how the fine-grained dynamics of HGB neural oscillations in hierarchically organized human motor regions predict movement speed during movement preparation and execution. More precisely, it is unclear how the transition from motor planning in the higher-order motor cortices to motor command in M1 affects movement speed in the human motor system. Furthermore, the primary and the non-primary motor cortices collaborate to achieve smooth movements. For example, the non-primary motor cortices are involved in the initiation of hand movements by modulating the primary motor region (Liu et al. 2002), and both contralateral and ipsilateral PM and SMA influence M1 during hand movements (Civardi et al. 2001; Buch et al. 2010; Groppa et al. 2012; Quessy et al. 2016). Here, it remains to be further elucidated how the communication (e.g., gamma-band coherence) between the PM, SMA, and M1 during movement planning and execution affects movement speed.

Given the extensive utilization of reaction times (RTs) as an indicator of movement preparation and execution, we employed the RTs in a cued finger movement task in the present study as a behavioural measurement of movement speed (Churchland, Santhanam*, et al.* 2006; Churchland, Yu*, et al.* 2006; Afshar *et al*. 2011; Kaufman et al. 2016; Paraskevopoulou et al. 2021). Taking advantage of the high temporal resolution of intracranial electroencephalography (iEEG) in patients with drug-resistant epilepsy and electrode coverage of the human PM, SMA, and M1, we investigated regional variations in peak amplitude, peak latency, and duration of HGB oscillations from motor planning to execution and how they correlate with variance in RTs. Although our results show significant activation in both the classical movement-related gamma and beta frequency bands, we focus on summarizing the neural activity characteristics of the gamma frequency band due to its close relationship with movement execution and preparation (Gonzalez et al., 2006; Keeley et al. 2019). In the current study, only HGB oscillations were foucused. Moreover, we investigated how the sequential activation and coherence of motor regions affected RTs. Voluntarily participating drug-resistant epilepsy patients with electrode coverage in PM, SMA, and M1 were asked to perform visually or auditorily cued left or right index finger movements. We predicted that the peak latency, peak amplitude, and duration of HGB activity in the motor system should correlate with the variance in RTs. Specifically, HGB activation should peak earlier (i.e., during the motor planning phase) in the higher-order motor regions (PM and SMA) and later (i.e., during the motor command phase) in the primary motor cortex (M1). Additionally, we further investigated how the sequential transition in HGB activation from motor planning in PM/SMA to motor command in M1 and the HGB phase coherence between PM/SMA and M1 contributed to reaction times.

## Materials and Methods

### Participants

Data were collected from 13 patients undergoing neurosurgical treatment for intractable epilepsy in Guangdong Sanjiu Brain Hospital, China. All electrodes were implanted stereotactically within one or both hemispheres (Fig. 1C). The placement of the electrodes, which was entirely based on clinical evaluation, was naive to the purpose of the present study. All patients had normal or corrected-to-normal vision (see Table 1 for demographic details), and no structural abnormalities, head trauma, or encephalitis were reported. The patients provided informed approval before the experiment. Given the absence of evoked motor-related responses in four participants’ implanted motor cortex electrodes, data from the four patients were excluded from detailed analyses (please see details in the ‘Electrode Reconstruction and Locations’ section). The School of Psychology, South China Normal University’s ethics committee, approved the study.

**Fig. 1.**
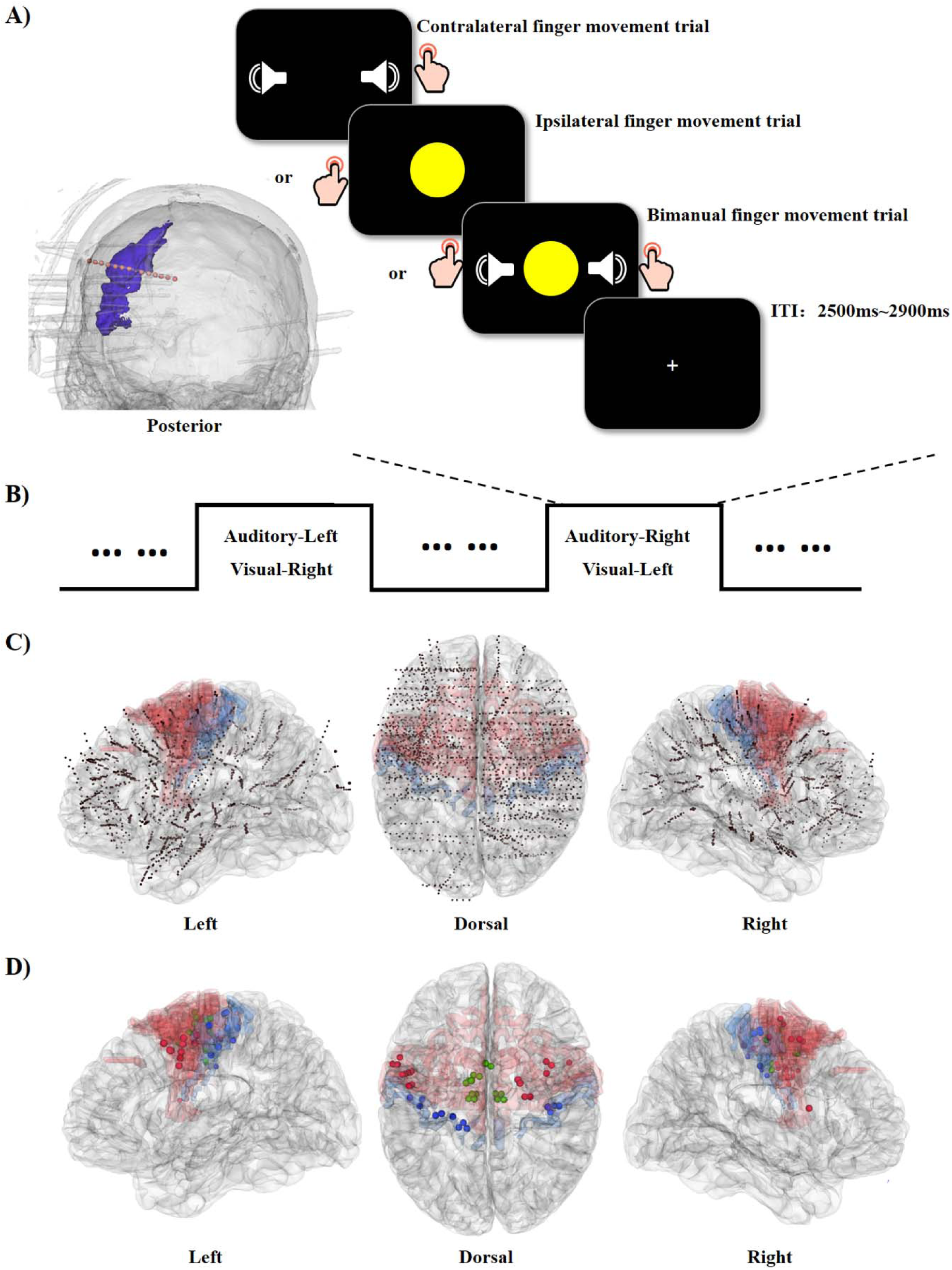
Paradigm and electrode positions of the three motor areas. **A)** Left: an example of how the stereo electrodes (red dots) penetrate the motor areas. The purple area is the reconstructed left precentral gyrus in one representative patient. Right: experimental paradigm. For example, in the “Auditory-right_Visual-left” blocks, the patients were instructed to move the right finger if they heard the auditory target while moving the left finger if they saw the visual cue and moving both fingers for the bimodal cues. Therefore, the finger movements in the unimodal auditory trials were contralateral while ipsilateral in the unimodal visual trials to the implanted motor electrodes. **B)** There were two types of blocks: “Auditory-Left_Visual-Right” and “Auditory-Right_Visual-Left”. The bottom-up inputs were identical, while the finger-modality mapping was reversed in the two types of blocks. **C)** Anatomical locations of all electrode contacts in the 13 patients, normalized to the MNI space. The contacts falling in Brodmann Areas 4 (blue shadow) and 6 (red shadow) were first anatomically selected. **D)** Subsequently, functionally responsive contacts within anatomically localized contacts were selected and categorized into the three motor areas: primary motor cortex (M1: blue electrodes), premotor cortex (PM: red electrodes), and supplementary motor area (SMA: green electrodes).

**Table 1.** Demographic information and electrode distribution in each patient.

| ID | Age/<br>Gender | Ictal or interictal | L/R<br>Handed | Epilepsy<br>duration | Task<br>blocks | M1<br>contacts | SMA<br>contacts | PM<br>contacts |
| --- | --- | --- | --- | --- | --- | --- | --- | --- |
| 1 | 22/F | Left insula | L | 10 years | 12 |  | 3 |  |
| 2 | 23/F | Left insula and temporal lobe | R | 17 years | 9 |  |  | 3 |
| 3 | 22/F | Left insula and inferior frontal | R | 17 years | 56 | 6 (1) | 4 | 6 |
| 4 | 26/M | Left inferior parietal | R | 12 years | 18 |  | 1 |  |
| 5 | 22/M | Left insula | R | 9 years | 52 | 2 (2) |  | 3 |
| 6 | **18/M** | **Right ACC** | **R** | **10 years** | **8** | **6 (1)** | **14 (2)** | **13 (3)** |
| 7 | **16/M** | **Left insula and frontal** | **L** | **1 year** | **25** | **4 (4)** | **10 (6)** | **22 (7)** |
| 8 | **24/M** | **Right precuneus and intraparietal sulcus** | **R** | **14 years** | **25** | **2 (1)** | **6 (2)** | **5 (2)** |
| 9 | 15/M | Right temporal and parietal | R | 10 years | 16 | 2 | 3 |  |
| 10 | 17/M | Left medial frontal | R | 14 years | 45 |  | 2 | 11 (3) |
| 11 | **21/M** | **Right parietal** | **R** | **3 years** | **44** | **2 (2)** | **2 (2)** | **2 (2)** |
| 12 | 23/M | Right frontal and insula | R | 17 years | 42 | 4 | 3 | 9 (1) |
| 13 | 24/M | Left superior prefrontal | R | 10 years | 12 | 4 (2) | 4 (3) | 5 |
| Total contacts and responsive contacts (in brackets) of each brain region |  |  |  |  |  | 32 (13) | 52 (15) | 79 (18) |

F: female. M: male. L: Left. R: Right. The number in brackets is the number of responsive electrodes. The patients in **bold** have responsive contacts simultaneously in M1, PM, and SMA. Due to the lack of responsive electrodes in patients 1, 2, 4, and 9, their data did not enter the group analysis.

### Stimuli and experimental design

Both visual and auditory cues were employed. The visual cue was a yellow-filled sphere presented centrally on a computer screen with a 1.5° visual angle radius. The auditory cue was a pure 4000 Hz tone. The visual and auditory cues were presented for 50 ms individually or simultaneously. A white cross (visual angle: 1° × 1°; RGB: 255, 255, 255) on a black background (RGB: 0, 0, 0) was used as the default visual display (Fig. 1A). The tasks were presented via Presentation Software (https://www.neurobs.com/) on a computer with a 23-inch screen (resolution, 1920 × 1080) positioned 60 cm from the patient’s eye level. The auditory stimuli were broadcasted by loudspeakers placed behind the computer monitor.

The following three trial types were presented: 1) only an auditory cue was presented for 50 ms; 2) only a visual cue was presented for 50 ms; and 3) the visual and auditory cues were presented simultaneously for 50 ms. The three trial types were presented randomly within one block. The experimental task consisted of i) pressing a button with the left index finger if the visual target was presented, ii) pressing a button with the right index finger if only the auditory target was presented, and iii) pressing both buttons with their left and right index fingers if the visual and auditory cues were presented simultaneously. The corresponding relationship between the finger and target was counterbalanced between the blocks (Fig. 1B). At the beginning of each block, patients were instructed on the correspondence between the target and the respective finger. The tasks for the participants were unilateral left finger movement, unilateral right finger movement and bilateral finger movement. However, only the unilateral left/right finger movement trials were submitted for further data analysis. The bilateral finger movement trials were used to select the responsive electrodes of the participant. For all unilateral finger movement trials, the definition of contralateral and ipsilateral movement was based on the relative position of the electrode hemisphere and the moving finger hemisphere. The moving finger was opposite to the hemisphere in which the electrodes were implanted for the contralateral finger movement condition. The moving finger and the electrodes’ hemisphere were on the same side for the ipsilateral finger movement condition. For instance, consider the seventh contact of one electrode implanted in one patient’s *left* primary motor cortex (Fig. 1A). In the Auditory-right_Visual-left block, the task of the patient was to press the button with the right index finger if he/she heard the auditory cue (in this case, it was the contralateral finger movement condition) and press the button with the left index finger if he/she saw the visual cue (in this case, it was the ipsilateral finger movement condition). The patient was supposed to press the buttons with both index fingers if both cues were presented simultaneously (bimanual movement condition). The intertrial intervals ranged from 2500 ms to 2900 ms (ITI: 2500 ms, 2600 ms, 2700 ms, 2800 ms, and 2900 ms) and were presented randomly.

All other trials were separated into two blocks: Auditory-Left_Visual-Right and Auditory-Right_Visual-Left. The Auditory-Left_Visual-Right and Auditory-Right_Visual-Left blocks were presented in alternating ways. The specific number of blocks for each patient was determined by the available research recording time in the clinical environment, which varied among patients (Table 1). Each block contained 50 trials, with the proportion of left finger movements, right finger movements, and bimanual movements kept at 2:2:1. Because the bimanual movement condition was used to localize the electrodes related to finger movement, fewer trials were included here to shorten the length of the experiment. The total number of trials varied among each participant. We randomly selected 150 trials for each condition from the overall dataset to avoid significant differences arising from the disparate trial counts. In the Auditory-Left_Visual-Right block, auditory targets were related to the left-finger movement, and visual targets were related to the right-finger movement. The relationship between target and finger movement was reversed in the Auditory-Right_Visual-Left block. Before the formal experiment, a 10-minute training test was presented to the participants to ensure that they understood the tasks and experimental procedures.

All trials were separated into (i) contralateral finger movements, (ii) ipsilateral finger movements, and (iii) bimanual movements. According to the reaction times, trials were further separated into the contralateral fast condition (contralateral fast), the contralateral slow condition (contralateral slow), the ipsilateral fast condition (ipsilateral fast), and the ipsilateral slow condition (ipsilateral slow). Fast conditions contained 30% fastest trials, and slow conditions contained 30% of the slowest trials.

### Intracranial EEG acquisition and preprocessing

Intracranial EEG data were acquired by the Nihon Kohden (Tokyo, Japan) monitoring system with a 1000 Hz sampling rate and a bandpass filter of 0.1–300 Hz at the patient’s bedside in Guangzhou Sanjiu Brain Hospital, China. All patients were implanted with depth electrodes containing 10-16 contacts (0.8 mm diameter and 2 mm width) with an intercontact spacing of 1.5 mm. To ensure good data quality, the contacts were only considered if no seizure was observed at least 1 hour before and after the experiments. The epileptic electrode selection and the artifact rejection of the epileptic spikes were instructed by a neurologist offline after the experiment. All data clips underwent trial-by-trial artifact rejection to eliminate any pathological activity, e.g., interictal epileptiform discharge (sFig. 1). Due to strict control of the participants’ state during data collection, the occurrence rate of interictal epileptiform discharges (IEDs) in the experimental data was minimal. A total of 5.4% of the trials showed IEDs in the contralateral condition, and 6.1% of the trials exhibited IEDs in the ipsilateral condition. Trials with IEDs were excluded from further analysis.

Signals were downsampled to 500 Hz and referenced with a bipolar montage. A 47-53 Hz notch filter was used to eliminate the line noise. All data analyses were conducted using Fieldtrip (http://www.fieldtriptoolbox.org/) functions and custom scripts in MATLAB 2016a (https://www.mathworks.com/products/MATLAB.html). Time-frequency decomposition was performed using the Morlet wavelet transformation function implemented in Fieldtrip to compute the power spectra of the electrophysiological signals. The time-frequency analysis was performed for each epoch with a frequency step of 1 Hz and a time window step of 1 ms. The time-frequency analysis focused on 2-140 Hz with different cycles for each frequency (the number of cycles linearly increased with the frequency from 2 to 30 cycles). Each epoch was baseline-corrected in decibels relative to a time window of 500 ms to 100 ms before the stimuli onset for each frequency band (dB power = 10 × log10(power/mean power of baseline)) to normalize the distributions of the power spectra for both time-locked on cue and time-locked on response. The power spectra were averaged for the HGB of 60-140 Hz. An extended epoch was employed in this study to mitigate the edge effects of the wavelet time-frequency analysis. Continuous intracranial EEG data were segmented into epochs ranging from 1500 ms before to 1500 ms after stimulus onset. The data containing the edges were excluded from further analyses. The time window subjected to statistical analysis spanned from the cue onset to 1000 ms after the cue onset with time 0 locked to the cue. For analyses with time 0 locked on the response, the time window spanned from −500 ms before to 500 ms after the response.

### Electrode reconstruction and locations

To identify the anatomical location of the electrodes, post-implantation computer tomography (post-CT) was co-registered with pre-implantation T1-weighted MR images using rigid affine transformations of the FLIRT algorithm in FSL (Jenkinson and Smith 2001). The co-registered images were visually inspected afterwards to guarantee adequate registrations. For each patient, the T1 image was segmented with FreeSurfer (Fischl et al. 1999) to obtain the Brodmann landmark of each brain region. The Brodmann landmarks were overlaid on the patient’s brain (Fig. 1C and D), and the structural location of each electrode was visually inspected by a medical doctor using 3D Slicer (https:www.slicer.org). The electrodes implanted in Brodmann Area 4 were defined as primary motor cortex (M1) electrodes. Electrodes in the lateral Brodmann Area 6 were the premotor cortex (PM) electrodes, and those in the medial wall of Brodmann Area 6 were the supplementary motor area (SMA) electrodes (Woolsey et al. 1950) (Fig. 1D, M1 electrodes in blue, PM electrodes in red, SMA electrodes in green).

Not all the electrodes anatomically located in the motor cortex are responsive to finger movements. Participants were cued to make either unilateral or bilateral finger movements. Since we were interested only in unilateral movements in the present study, only the unilateral finger movement (either contralateral or ipsilateral to the side of the implanted electrodes) trials were included in the current analyses. The bilateral finger movement trials, on the other hand, were used as a group of independent trials to screen the motor cortex electrodes responsive to finger movements. Gamma activity increases and beta activity decreases in the motor system have been well documented during movement planning and execution (Cheyne and Ferrari 2013; Kilavik et al. 2013; Tatti et al. 2022). Therefore, the electrodes that anatomically sit in the motor system and functionally showed gamma power increases and beta power decreases during the bilateral finger movement trials were selected as the functioning motor electrodes and used for the analyses in the unilateral finger movement trials (sFig. 2). If none of the motor electrodes were responsive in the bimanual movement trials of a patient, the patient was excluded from the analysis. Therefore, data from nine patients were submitted for further analyses (Fig. 1D). For the 9 patients, there were 18 responsive electrodes in the PM, 15 in the SMA, and 13 in the M1 regions.

**Fig. 2.**
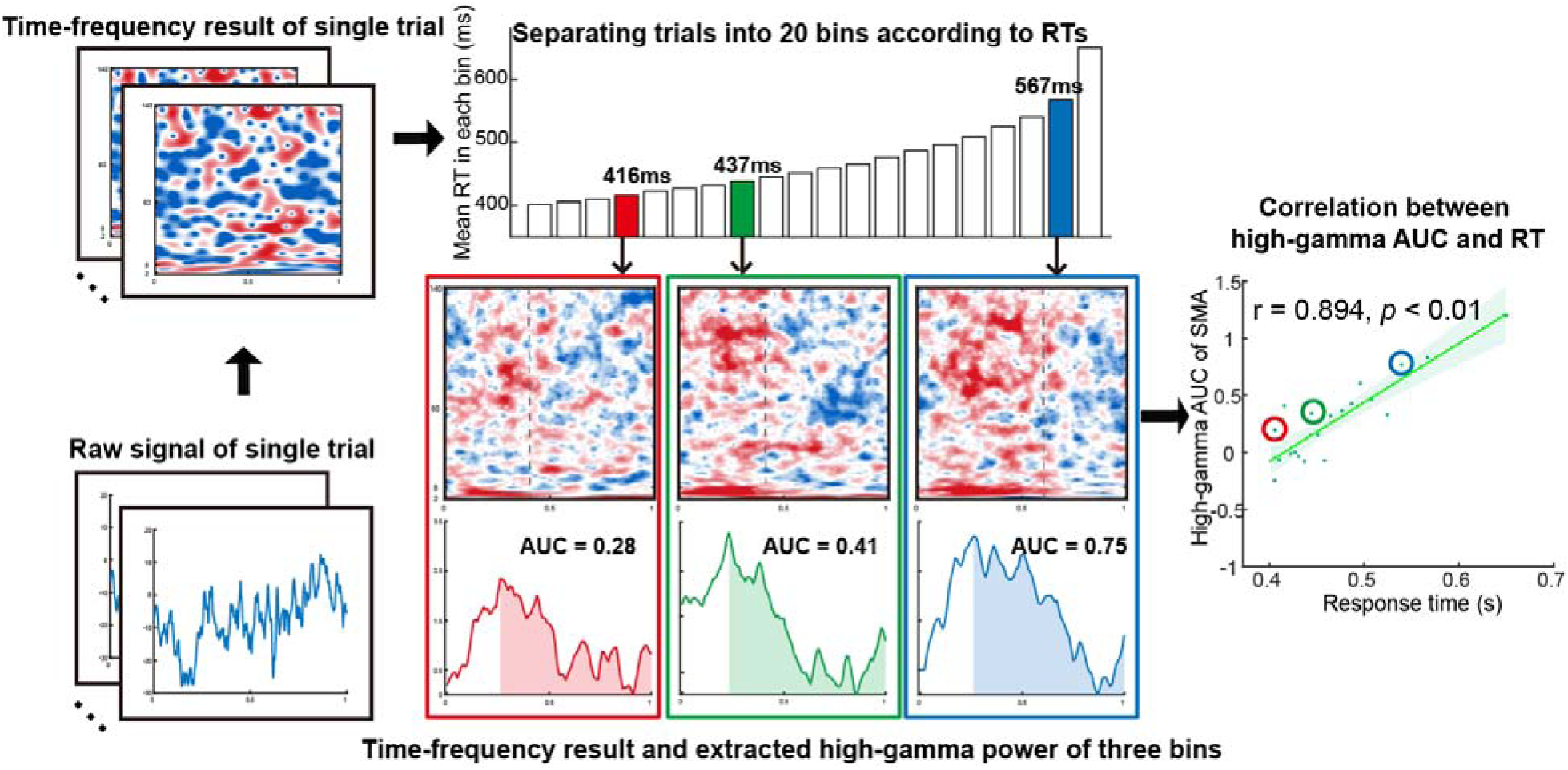
Linear mixed model analysis between high-gamma AUC and RT. Schematic illustration of the procedure to calculate the correlation between the area under the gamma power curve (AUC) and the RTs. First, we performed a time-frequency analysis for each trial (the left upper panel). Second, all the trials were sorted by the RTs and separated into 20 consecutive bins (the middle upper panel). Within each trial bin, the power results and the RTs were averaged. For example, the mean RT of the fourth bin is 416 ms (the middle upper panel in red). The power of the averaged high-gamma band (60-140 Hz) within each trial bin was then extracted (time window: from the onset of the cue to 1000 ms after the cue onset). For example, in the middle lower panel, the red line represents the average power for the fourth trial bin, the green line for the eighth bin, and the blue line for the nineteenth bin. Finally, the high-gamma AUC was extracted, submitted to the linear mixed model, and correlated with the mean RT across all bins (the right panel: the dots in the red/green/blue circle indicate the data in the fourth\eighth\nineteenth bin). Shaded regions in the middle lower panels identify the high-gamma AUC in a given trial bin. The AUC represents sustained power activation from the peak to the end of the trial. The high-gamma AUC and RT correlation defines how sustained gamma activation affects finger movements. The grey dashed line shows the averaged reaction time in each trial bin.

For demonstration, the coordinates of the electrodes of each patient were normalized to the Montreal Neurologic Institute (MNI) space and visualized using 3D Slicer software (Fig. 1C). The MNI coordinates of all electrodes were projected to fsaverage standard space only for visualization purposes (Fig. 1C and D).

### Analysis of behavioral data

Behaviorally missed and incorrect trials were excluded from further analysis for all trials. The correct contralateral and ipsilateral trials were then separated into fast and slow trials per reaction time. Mean RTs were then submitted to a 2 (response hand relative to electrode sites: contralateral vs. ipsilateral) × 2 (response speed: fast vs. slow) repeated-measures ANOVA. Planned paired *t* tests (with Bonferroni correction) were used to test the simple effects.

### Analysis of neural data

#### Contralateral movement effect in the motor cortex

Our study aimed to investigate the HGB (60-140 Hz) signal dynamics of iEEG signals in motor areas (M1, PM, and SMA). The HGB power was extracted and averaged to compare the difference between conditions. A non-parametric permutation test was used to control the multiple-comparisons problem. To avoid double dipping (Kriegeskorte et al. 2009), no statistical tests were performed on the full power spectrum of 2-140 Hz.

Here, we defined the contralateral movement effect as significantly higher activation of the HGB power in the contralateral than in the ipsilateral condition. The HGB power of the contralateral and ipsilateral trials was extracted and averaged for M1, PM, and SMA to test the contralateral movement effect. We then compared the averaged HGB power of the contralateral vs. ipsilateral conditions using a non-parametric permutation test with correction for multiple comparisons across electrodes for the three motor areas.

#### The relationship between HGB power in the motor cortex and reaction time

Trials were separated into fast and slow trials per reaction time to explain the relationship between the motor cortex’s HGB power and RT. The fastest 30% of trials were defined as fast trials, and the slowest 30% of trials were defined as slow trials. The HGB power was extracted and averaged from the contralateral-fast, contralateral-slow, ipsilateral-fast, and ipsilateral-slow conditions for M1, PM, and SMA. Within each motor area, the averaged HGB power of the fast vs. slow trials was compared for the contralateral and ipsilateral conditions, respectively, i.e., “contralateral-fast vs. contralateral-slow” and “ipsilateral-fast vs. ipsilateral-slow”, using a non-parametric permutation test with correction for multiple comparisons across electrodes.

The time interval from the stimulus onset to the time point of maximum HGB power, i.e., the peak, was defined as the peak latency of the fast and slow trials. The maximum HGB power amplitude was defined as the peak amplitude of the fast and slow trials. The peak latency and amplitude were extracted and submitted to paired *t* tests to compare the difference between fast and slow trials in the contralateral and ipsilateral conditions, respectively, for all three motor areas.

The linear mixed model (LMM) was applied to reveal a possible relationship between the HGB power of the motor cortex and reaction times. A trial-by-trial correlation between the HGB power and reaction times was tried first. However, since the validity of detecting the peak power of a single trial is severely compromised by non-specific noises (sFig. 3), we chose the method of trial bins to increase the signal-to-noise ratio. Specifically, all trials were divided into 20 bins (from 400 ms to 800 ms, with a step of 20 ms, Fig. 2, middle upper panel). For each bin, the HGB power and reaction time were extracted and averaged (Fig. 2, middle lower panel, the red\green\blue lines indicate the averaged HGB power in the fourth\eighth\nineteenth bin, respectively). The peak activation of gamma power was defined as the local maximum activation of the extracted gamma power within the time window from cue stimulation onset to 1000 ms after the cue. The peak latency represented the period from the cue onset to the timestamp of the peak activation. The area under the curve referred to the region between the peak activation and 1000 ms after the cue (Fig. 2, middle lower panel, red/green/blue shaded areas). The calculation of the LMM was based on the averaged RTs and averaged HGB power of 20 bins for each electrode.

**Fig. 3.**
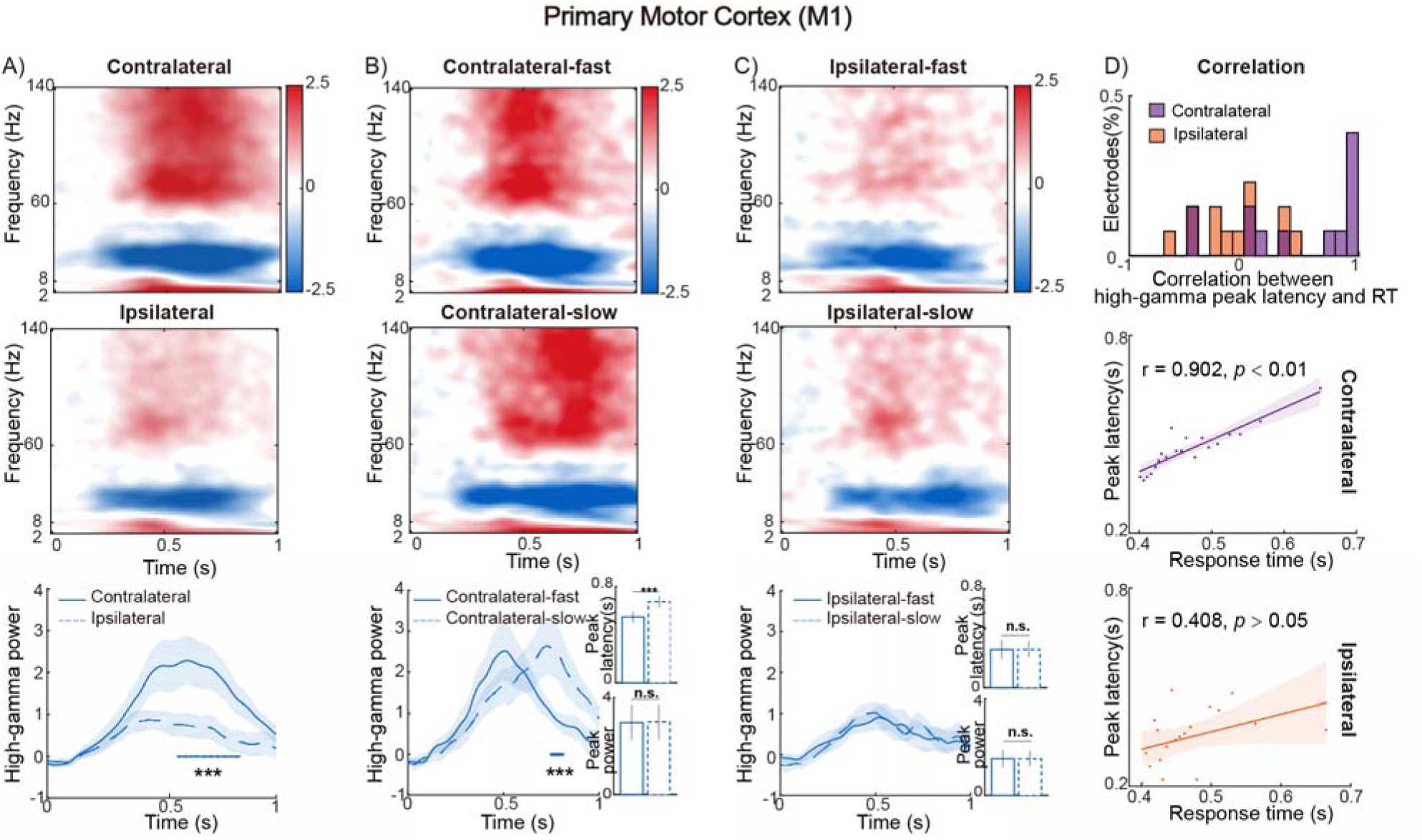
Time-frequency results in M1. **A)** Group-level time-frequency results of the primary motor cortex for the contralateral (upper panel) and ipsilateral (middle panel) movement conditions. Time ticket zero marks the onset of the stimulus. The temporal profiles of the high-gamma power (60-140 Hz, the lower panel) are displayed as a function of the contralateral (blue solid line) vs. ipsilateral (blue dashed line) trials. **B)** Time-frequency spectra in the contralateral movement condition as a function of the fast (upper panel) vs. slow (lower panel) contralateral finger movements. **C)** Time-frequency spectra in the ipsilateral movement condition as a function of the fast (upper panel) vs. slow (lower panel) ipsilateral finger movements. In **B)** and **C)**, the temporal profiles of high-gamma power (lower panels) are displayed as a function of the fast (solid blue lines) vs. slow (dashed blue lines) trials for the contralateral and ipsilateral movements, respectively. The peak latency and the peak amplitude of the high-gamma power were extracted and submitted to paired *t* tests between the fast and slow trials for the contralateral and ipsilateral movements, respectively (the inserted bar figures). Error bars represent the standard error of the mean (SEM). **D)** Distribution of all the responsive M1 electrodes as a function of the correlation value between the high-gamma power peak latency and the RTs in the contralateral (upper panel in purple) and ipsilateral (upper panel in orange) movement conditions. The correlation between the high-gamma power peak latency and the RT in the same representative M1 electrode in the contralateral (middle panel) and ipsilateral (lower panel) movement conditions is shown in the scatter figure. *** *p* < 0.01; non-parametric permutation test with correction for multiple comparisons across electrodes. All plot shadows around the solid/dashed lines indicate the SEM.

The LMM analysis was performed in two ways. First, the electrodes were entered as a random factor, the peak latency in the bin power (i.e., the maximum activation within the averaged HGB power) was entered as a fixed factor, and the averaged RTs of each bin were entered as a dependent factor. While controlling for the variability of the electrodes, the LMM calculation between the HGB peak latency and the RT indicated a relationship between the maximum activation of power and the behavioural response. Second, the area under the curve from the peak of HGB power to the end of the trial was extracted (Fig. 2, shaded regions in the middle lower panel) and used as a fixed factor in the LMM. This area under the power curve defined how sustainably the power was enhanced from the maximum activation to the end of the trial. The LMM calculation between this area and the average RT was computed to investigate a possible relationship between the sustained activation of the power and the reaction time (Fig. 2, right panel). For demonstration, the correlation coefficients between the HGB power peak latency/the area under the curve and the RTs were computed for each electrode based on the bin data.

#### Prediction of how non-primary motor activity affects primary motor activity

We then determined whether there was evidence for precise timing of motor-related HGB responses, differentiating the non-primary and primary motor cortex during finger movements. For the motor-related analysis, we focused on the contralateral movement conditions. To ensure that differences in power were not affected by individual patients, we also selected four patients with electrodes in all three motor regions (Table 1 in **bold**). HGB power of the three motor areas in the contralateral condition was extracted and averaged. The peak latency across electrodes of the three brain regions was calculated and submitted to a one-way ANOVA to compare the HGB peak latency between M1 and PM/SMA. Post hoc comparison of peak latencies between two brain regions with Bonferroni correction for multiple testing was performed. To better demonstrate the motor command function of M1, trials were re-epoched and time-locked to the response onset rather than the cue onset. All the other time-frequency decomposition procedures were kept the same as mentioned before.

Last, to further investigate the relationship between the peak latency in the M1 region and the sustained activation of the PM and SMA regions, the linear mixed model (LMM) was applied. The electrodes were entered as a random factor, the peak latency of M1 was entered as a fixed factor, and the area under the curve of PM and SMA were entered as dependent factors. This LMM model depicted how the sustained activation of the non-primary motor cortex affected the motor command of the primary motor cortex.

#### Pairwise phase consistency between the primary and non-primary motor cortex for fast and slow trials

Pairwise phase consistency (PPC) was calculated to analyse phase consistency across trials between the different motor areas. The PPC gives an unbiased estimate of the population statistics of the squared phase-locking value (PLV) (Vinck et al. 2010; Rohenkohl et al. 2018). The PPC was computed from the cross-spectral densities (CSDs) obtained from the wavelet convolution described in the abovementioned time-frequency analysis section. The CSDs were used to find mutual resonant frequencies in a pair of signals. The CSDs were computed as dB change relative to the same baseline of 500 ms to 100 ms before stimuli onset. The phases in the CSDs represent the phase differences between the oscillations of the electrode pairs. The PPC was calculated as follows:

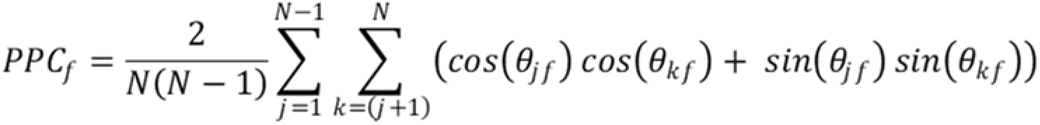

where *j* = (1, 2, …, *N*), *N* is the number of all observations (trials), *θ_j_* and *θ_k_* are the relative phases between two signals in a particular frequency band, and *f* is the specific frequency of HGB (60-140 Hz). The PPC was calculated between electrode pairs of PM-M1, SMA-M1, and PM-SMA.

### Statistical analysis

The number of patients was limited in our study. Therefore, the group and statistical analyses were based on the electrode level rather than the patient level. However, since the electrode number from each patient was not always the same, a weighted procedure was adopted to ensure that each patient contributed equally to the statistics (The weight equals the number of electrodes for the current patient divided by the total number of electrodes).

To calculate the significance of power and phase consistency differences between conditions, high-gamma band (HGB, 60-140 Hz) data were extracted, averaged, and statistically tested using a non-parametric permutation test with correction for multiple comparisons across electrodes (Maris and Oostenveld 2007). For instance, when calculating the difference in high-gamma power between the contralateral and ipsilateral finger movement conditions, we first performed paired *t* tests at each time point. A time point that passed a threshold *p* value smaller than 0.01 was marked as significant. The multiple comparisons were based on considering the cluster as the unit for determining the threshold (a cluster was defined as a group of contiguous suprathreshold time points). The total *t* value within a cluster represented the real *t* value of this cluster. Then, a cluster-based correction was adopted to define whether the real *t* value was significant. The data of both conditions were randomly shuffled 1000 times, and for each shuffle, the total *t* value of the largest significant cluster was entered into the distribution of the null hypothesis. The real data cluster was considered significant if the real cluster *t* value exceeded 99% of the null distribution at α = 0.01.

The linear mixed model (LMM) was adopted to investigate the relationship between the HGB neural response of the motor areas and the behavioural response. The LMM approach allowed us to adjust for confounding factors by considering the contributions of individual electrodes (Baayen et al. 2008; Bates et al. 2015; Kuznetsova et al. 2017). The packages implemented in R were adopted to construct the LMM model and calculate the significance (lme4 and lmerTest package; lme4: construct LMM model; ImerTest: obtained the *p* values of the LMM model). The neural responses and the reaction times were fitted with a random intercept model, and the LMM coefficients represented a correlation between them. The significance level was set at α = 0.01.

## Results

To better understand the relationship between reaction times and the underlying neural dynamics in the motor regions, the contralateral and ipsilateral trials were categorized into fast vs. slow trials (see Methods). The mean RTs in the resulting four conditions were then submitted to a 2 (side of response hand: contralateral vs. ipsilateral) × 2 (response speed: fast vs. slow) repeated-measures ANOVA (sFig. 4). The main effect of response speed was the only significant effect, *F* _(1, 8)_ = 286.243, *p* < 0.01. Neither the main effect of the response hand nor the interaction was significant, both *p* > 0.01, indicating that the contralateral and ipsilateral RTs were comparable.

**Fig. 4.**
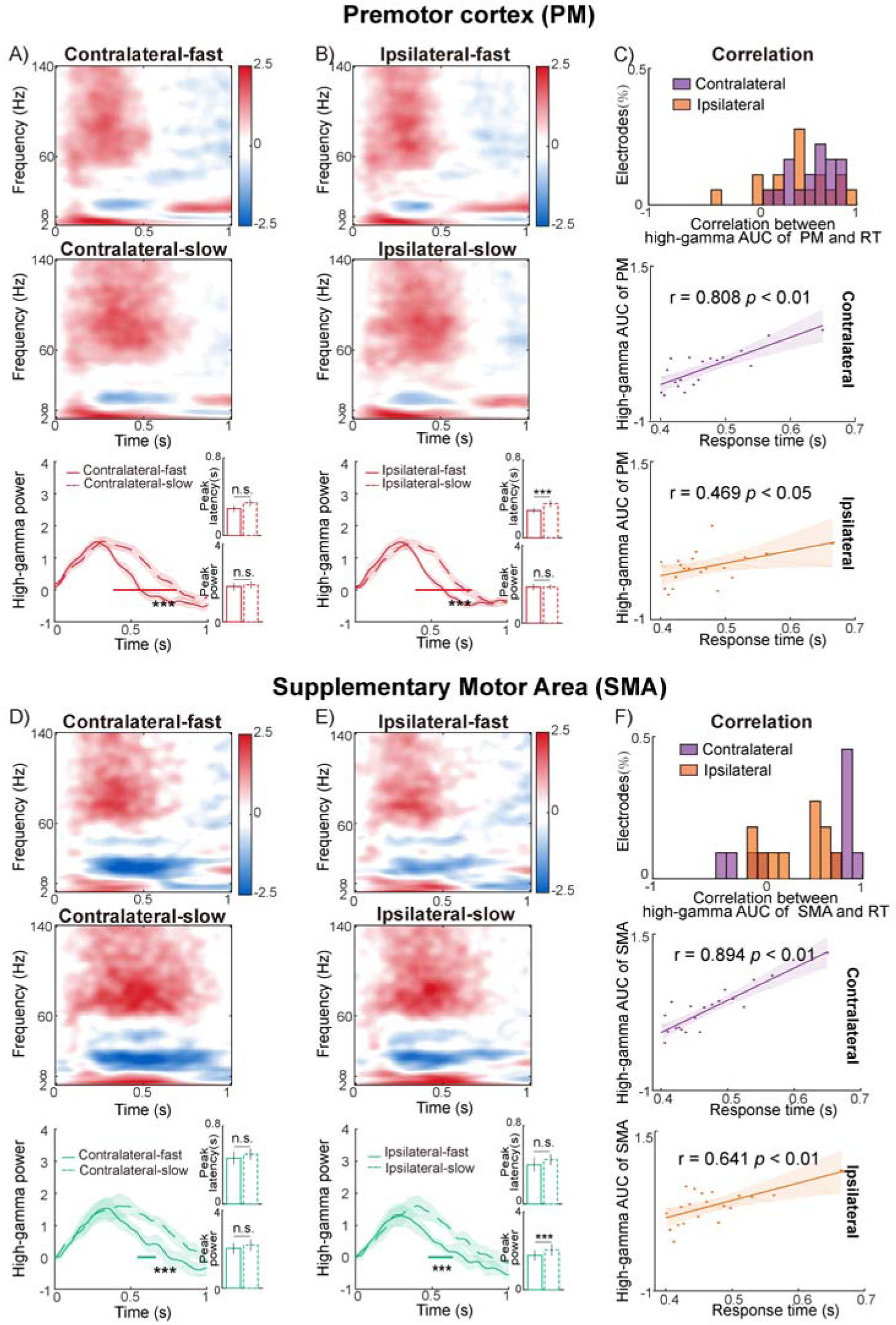
Time-frequency results of PM and SMA for fast and slow trials. **A)** Group-level time-frequency results of contralateral movements within the premotor cortex for fast trials (contralateral-fast, upper panel) and slow trials (contralateral-slow, middle panel). Time ticket zero marks the onset of the stimulus. The temporal profiles of the high-gamma power (60-140 Hz, lower panel) are shown as a function of the contralateral-fast vs. contralateral-slow trials. The peak latency and the peak amplitude of high-gamma power were extracted and submitted to paired *t* tests (the inserted bar figures). Error bars represent the standard error of the mean (SEM). **B), D), and E)** are organized similarly as in **A)**. **B)** shows the results of the PM region in the ipsilateral movement conditions, **D)** shows the results of the SMA region in the contralateral movement conditions, and **E)** shows the results of the SMA region in the ipsilateral movement conditions. **C)** Distribution of all the responsive PM electrodes as a function of the correlation value between the high-gamma area under the curve (AUC) and the RT in the contralateral movement conditions (upper panel in purple) and ipsilateral movement conditions (upper panel in orange). The correlation between the high-gamma AUC and the RT in the representative electrode of PM in the contralateral movement conditions (middle panel) and ipsilateral movement conditions (lower panel) is shown in the scatter figure. **F)** The same as **C)**, except for the SMA region. *** *p* < 0.01; non-parametric permutation test with correction for multiple comparisons across electrodes. In **A) B) D)**, and **E)**, the shadows around the solid/dashed lines indicate the SEM. The shadow around the lines in **C)** and **F)** indicates 95% confidence intervals around the linear fit line.

### Contralateral finger movement effect in M1

During both contralateral and ipsilateral finger movements, M1, PM, and SMA all showed basic motor-related effects, i.e., event-related synchronization (ERS) in the HGB and event-related desynchronization (ERD) in the alpha-beta bands (Fig. 3 and 4, sFig. 5) (Crone et al. 1998, Miller et al 2007). The HGB ERS peaked at approximately 0.58±0.04 s (mean ± sem) in M1 (Fig. 3A) and at approximately 0.31 ± 0.03 s and 0.44± 0.05 s in SMA and PM, respectively (sFig. 5). Furthermore, only in the M1 region was the amplitude of the HGB power significantly higher in the contralateral than in the ipsilateral condition, *p* < 0.01, non-parametric permutation test with correction for multiple comparisons, i.e., a significant contralateral finger movement effect was present (Fig 3A lower panel). However, the HGB power in the PM and SMA was equivalently activated during contralateral and ipsilateral finger movements (sFig. 5A and B, right panel).

**Fig. 5.**
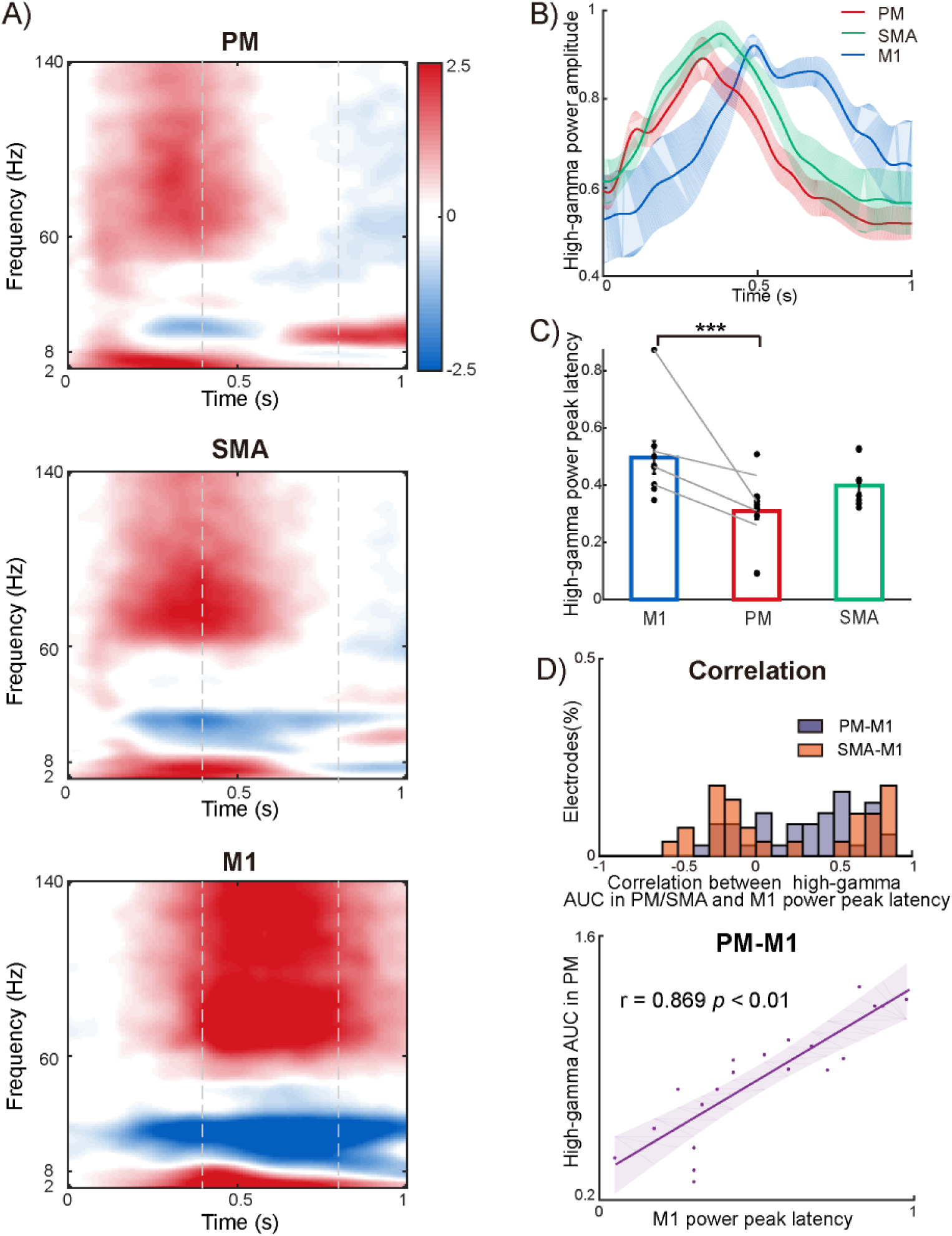
Sequential activation of PM, SMA, and M1. **A)** The high-gamma power activated sequentially across PM, SMA, and M1 for the contralateral movement condition. The grey dashed lines represent the RT range from the fastest (left) to the slowest (right). **B)** The mean time courses of the high-gamma power were extracted from PM, SMA, and M1. The shadow around each line indicates the standard error of the mean (SEM). **C)** The peak latency of the high-gamma power in the electrodes of three motor areas was submitted to a one-way ANOVA. Each black dot represents one contact. Each grey line connects the mean peak latency of all electrodes in the M1 vs. the PM area in one patient. **D)** Distribution of all the responsive PM/SMA and M1 electrode pairs as a function of the correlation value between the high-gamma AUC of PM (upper panel in purple)/SMA (upper panel in orange) and the power peak latency of M1 in the contralateral movement conditions. The correlation between the high-gamma AUC of PM (lower panel) and the power peak latency of M1 in a representative electrode pair in the contralateral movement conditions is shown in the scatter figure. All these results were based on four patients with electrodes implanted simultaneously in the PM, SMA, and M1. *** p < 0.01; with Bonferroni correction. The shadow around the line in **B)** and error bars in **C)** indicate SEM. The shadow around the line in **D)** indicates 95% confidence intervals around the linear fit line.

When the trials were split into fast vs. slow trials, the peak latency of HGB power in M1 was significantly shorter in the fast than slow trials, only in the contralateral finger movement condition, *p* < 0.01, non-parametric permutation test with correction for multiple comparisons, rather than in the ipsilateral movement condition (Fig. 3C lower panel). To further verify whether the peak latency of HGB power correlated with RTs on a trial-by-trial basis, the linear mixed model (LMM) coefficient was calculated. In the contralateral condition, the LMM showed a significant coefficient of *β* = 447.60, *p* < 0.01, for the peak latency of HGB power in M1 and the variance in RTs. To better understand the correlation between the peak latency of HGB and RTs, all trials were separated into 20 bins according to RTs, and the peak latency within each bin was extracted (see Methods). The correlation coefficient between the peak latency of HGB in M1 and RTs was calculated for every electrode within the trial-bin data. Among all M1 electrodes, 84.6% of the electrodes showed a positive correlation between the HGB peak latency and the RTs (Fig. 3D upper panel in purple). The correlation between the HGB peak latency and the RTs of a representative electrode is shown for demonstration purposes: the shorter the HGB peak latency, the faster the RT, r = 0.902, *p* < 0.01 (Fig. 3D middle panel). In the ipsilateral condition, the LMM showed no significant correlation between the HGB peak latency of M1 and the RT, *β* = 38.85, *p* > 0.01. Only 46.1% of the M1 electrodes showed a positive correlation between the HGB peak latency and the RT in the ipsilateral condition (Fig. 3D upper panel in orange).

### Prolonged HGB ERS in the PM and SMA impaired RTs

In the PM region, for both the contralateral and ipsilateral conditions, the HGB ERS lasted significantly longer in the slow trials than in the fast trials, *p* < 0.01, non-parametric permutation test with correction for multiple comparisons (Fig. 4A and B lower panels). We further adopted the linear mixed model (LMM) to evaluate how the duration of the HGB ERS in PM affected the RTs. Specifically, we calculated the LMM correlation between the area under the HGB power curve (AUC) and the RTs (Fig. 2, middle lower panel in shadow). In both the contralateral and ipsilateral conditions, the LMM revealed a significant coefficient between the HGB AUC and the RT, *β* = 151.74, *p* < 0.01 in the contralateral condition, and *β* = 159.09, *p* < 0.01 in the ipsilateral condition. Among all PM electrodes in all patients, 100% showed a positive correlation between the HGB AUC and the RT in the contralateral condition and 94.4% in the ipsilateral condition (Fig. 4C upper panel, contralateral condition in purple and ipsilateral condition in orange). For demonstration purposes, the correlation between the AUC of HGB and the RT is shown for one representative PM electrode for the contralateral and ipsilateral conditions, respectively, contralateral: r = 0.808, *p* < 0.01, ipsilateral: r = 0.469, *p* < 0.05 (Fig. 4C middle and lower panels). The SMA showed similar patterns as the PM. The HGB power activation lasted significantly longer in the slow trials than in the fast trials, *p* < 0.01, non-parametric permutation test with correction for multiple comparisons (Fig. 4D and E lower panels). Additionally, the LMM revealed a significant coefficient between the HGB AUC and RTs, *β* = 128.93, *p* < 0.01 in the contralateral condition and *β* = 94.85, *p* < 0.01 in the ipsilateral condition. Among all the SMA electrodes, 72.7% showed a positive correlation between the HGB AUC and the RT in the contralateral condition and 81.8% in the ipsilateral condition (Fig. 4F upper panel, contralateral condition in purple and ipsilateral condition in orange).

For demonstration purposes, the correlation between the AUC of HGB and the RT was shown for one representative SMA electrode for both the contralateral and ipsilateral conditions, contralateral: r = 0.894, *p* < 0.01, ipsilateral: r = 0.641, *p* < 0.01 (Fig. 4F middle and lower panels).

Taken together, prolonged HGB activation in the higher-order motor cortices, such as the PM and SMA, correlated with the RTs for both contralateral and ipsilateral finger movements: The slower the HGB power in the higher-order motor cortices returned to baseline, the longer the RTs.

In addition, for both the PM and SMA, the peak latency and peak amplitude of HGB were comparable between the fast and slow trials for contralateral finger movements (Fig. 4A lower right panel; Fig. 4D lower right panel). For the ipsilateral movements, however, the HGB peak latency in the PM was shorter in the fast trials than in the slow trials (Fig. 4B lower right panel), and the HGB peak amplitude in the SMA was higher in the slow trials than in the fast trials (Fig. 4E lower right panel).

### Prolonged HGB ERS in the PM predicted a delayed HGB power peak in M1 during contralateral movements

In four patients with implanted electrodes in all three motor regions, we found that the HGB ERS peaked sequentially in PM, SMA, and M1 during contralateral movements: earlier in PM (0.31±0.03 s) and SMA (0.39±0.03 s) and later in M1 (0.49±0.06 s) (Fig. 5A, 5B). When we time-locked the neural events to the response, it became evident that PM and SMA peaked before movement execution (sFig. 6A upper and middle panel, 6B), while M1 peaked at the onset of the movement (sFig. 6A lower panel, 6B). The peak latency of each electrode in the three regions was submitted to a one-way ANOVA followed by post hoc *t* tests with Bonferroni correction. We found a statistically significant difference in the HGB power peak latency between the three regions (*F*_(2, 30)_ = 6.828, *p* < 0.01). Post hoc *t* tests revealed a significantly earlier HGB peak in PM than in M1 (*t*_(20)_ = 3.329, *p* < 0.01, Fig. 5C). Similar temporal patterns were found when the neural events were time-locked to the response (sFig. 6C).

**Fig. 6.**
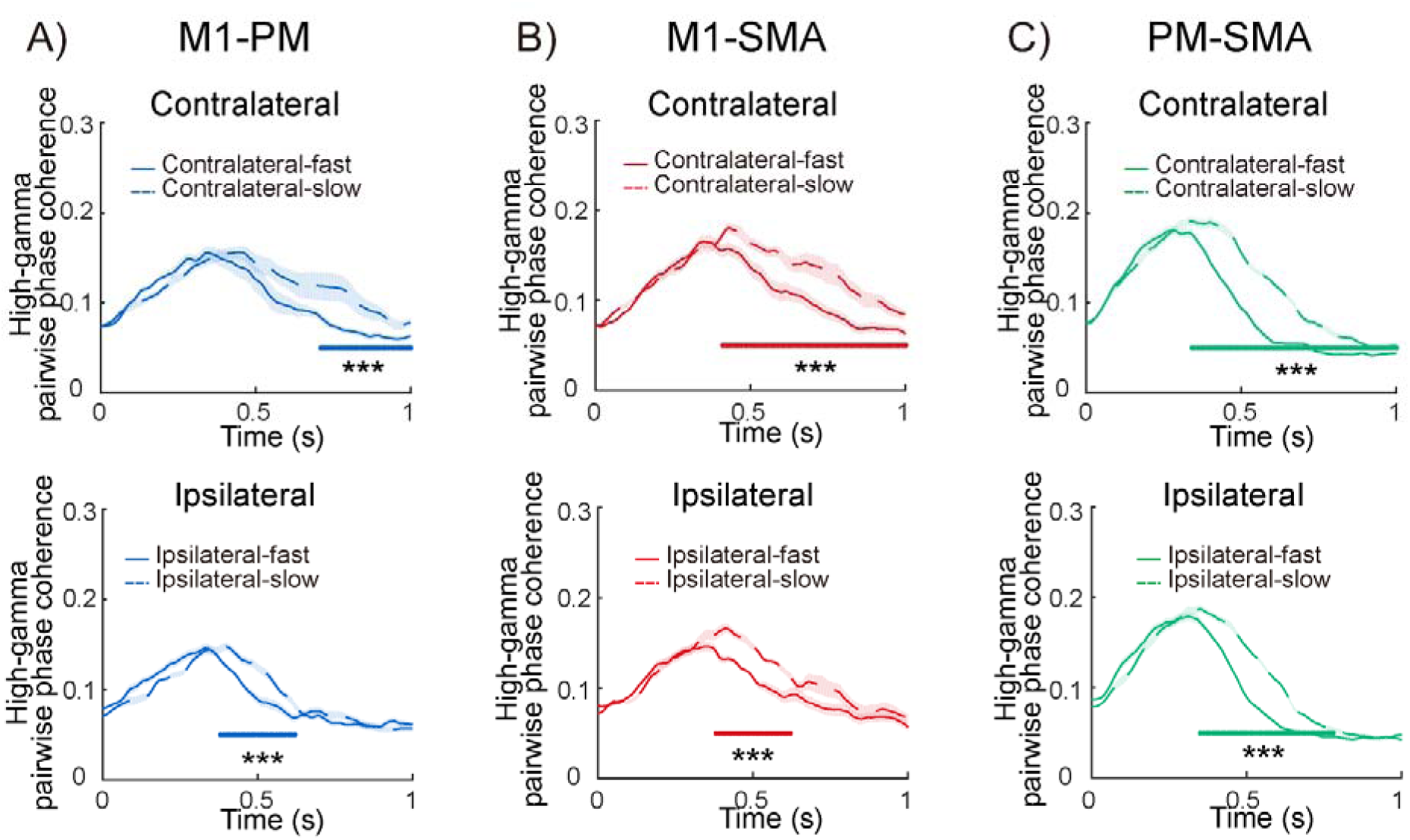
Pairwise phase consistency (PPC) results. **A)** PPC between PM and M1 as a function of the fast vs. slow trials in contralateral and ipsilateral movement conditions. **B)** PPC between the SMA and M1 for the fast vs. slow trials in the contralateral and ipsilateral movement conditions. **C)** PPC between the PM and SMA for the fast vs. slow trials in the contralateral vs. ipsilateral movement conditions. The extracted time courses of PPC in the high-gamma band were submitted to the non-parametric permutation test with correction for multiple comparisons across electrodes between the fast and slow trials. *** *p* < 0.01. The shadow around each line indicates the standard error of the mean.

The results within one region showed that both the peak latency of M1 (Fig. 3D) and the area under the curve (AUC) of PM and SMA (Fig. 4C and F) were highly correlated with the RT. In patients with simultaneous implantations in the non-primary and primary motor regions, we further examined whether the duration of the preparatory signals (i.e., the area under the curve, AUC) in the higher-order motor regions predicted the peak latency of the movement execution signals in M1. The LMM showed a significant correlation between the AUC of PM and the peak latency of M1, *β* = 0.123, *p* < 0.01. Among all the PM-M1 electrode pairs, 78.3% showed a positive correlation between the AUC of PM and the peak latency of M1 (Fig. 5D upper panel in purple). The correlation of a representative PM-M1 electrode pair is shown for demonstration purposes: the smaller the AUC of PM, the earlier the peak power latency of M1, r = 0.869, *p* < 0.01 (Fig. 5D lower panel). The LMM results on the SMA-M1 electrode pairs revealed no significant coefficient between the AUC of the SMA and the peak latency of M1, *β* = 0.017, *p* > 0.01. Only 50% of the SMA-M1 electrode pairs showed a positive correlation between the AUC of the SMA and the peak latency of M1 (Fig. 5D upper panel in orange).

Among the three regions in the motor system, the HGB ERS in the PM and SMA peaked earlier during motor preparation, while M1 peaked later around motor command. Moreover, the duration of the preparatory signals in PM predicted the latency of the motor command signals in M1.

### Prolonged HGB coherence between motor regions impaired RTs

We further calculated the pairwise HGB synchronization between the three regions, i.e., PM-M1, SMA-M1, and PM-SMA, for the fast and slow trials in the contralateral and ipsilateral conditions, respectively. For all three connections, the PPCs of the slow trials were more sustained than the PPCs of the fast trials for both contralateral and ipsilateral movements, *p* < 0.01, nonparametric permutation test with correction for multiple comparisons (Fig. 6A-C). The PPCs between the electrode pairs reflect the mean synchronization of all trials and thus cannot be interpreted on a single trial level. Accordingly, it is impossible to calculate the correlation between the PPCs and the RTs on a trial-by-trial basis.

## Discussion

In the present study, we delineated the fine-grained temporal dynamics of the high-gamma power in M1, PM, and SMA during motor planning and execution. Furthermore, we traced how the peak power, peak latency, duration of high-gamma power in M1, PM, and SMA, and interregional coherence correlated with the trial-by-trial variance in the speed of finger movements (i.e., RTs).

For the primary motor cortex (M1), the HGB power peaked approximately 400-800 ms after cue onset, and the HGB peak was strictly time-locked to movement execution (Fig. 3A, Fig. 5A lower panel, and sFig. 6A lower panel), indicating motor executive function. Accordingly, a significant contralateral finger movement effect was revealed in M1 (Fig. 3A lower panel). Evoked HGB activation in M1 is responsible for the execution of contralateral movements (Crone *et al*. 1998; Pfurtscheller *et al*. 2003; Brovelli et al. 2005; Szurhaj *et al*. 2005; Wiesman et al. 2021). For example, the maximum activation of HGB power in M1 is consistently centred on movement execution (Fonken *et al*. 2016). Moreover, the HGB power of M1 is evoked contralateral to the moving body parts and shows greater somatotopic organization than the lower frequency bands (Miller *et al*. 2007). Our results thus nicely replicated the contralateral movement function of the human primary motor cortex. More importantly, the peak latency of HGB power was found to correlate with the trial-by-trial variance in RTs of the contralateral finger movements in humans (Fig. 3B and D). The latter results suggest that the slower the motor command of the contralateral hand is, the later the M1 HGB power peak.

For the non-primary motor cortices (PM and SMA), the HGB power peaked approximately 300 ms after motor cue onset (Fig. 5A. upper and middle panel, sFig. 5A and B right and middle panel, sFig. 6A upper and middle panel), indicating an earlier motor planning function. For both the contralateral and ipsilateral finger movements, the duration of the HGB activation positively correlated with the RTs: the more sustained the HGB activation in the higher-order motor regions, the slower the finger movements were executed (Fig. 4A, B, D, and E lower panel). Alternatively, these results suggest that the duration of the preparatory motor planning signals in the PM and SMA could negatively influence the speed of subsequent finger movements (Fig. 4C and F). It has been well documented that the non-primary motor cortices in the PM and SMA are responsible for movement preparation before motor execution (Churchland, Yu*, et al.* 2006; Shibasaki and Hallett 2006; Churchland and Shenoy 2007; Miller *et al*. 2007; Nachev *et al*. 2008; Di Russo et al. 2017). For example, microstimulations in the PM region disrupt the preparatory activity of the PM region, resulting in a highly specific increase in RTs (Churchland and Shenoy 2007). The HGB power of the PM region peaks and prevails before movement execution and starts to return to baseline before the actual movement onset (Coon et al. 2016; Paraskevopoulou *et al*. 2021). Additionally, neurons in the SMA of monkeys show enhanced firing rates after the preparatory cue and before movement execution, indicating an essential role of the SMA during movement preparation (Nachev *et al*. 2008). The human SMA also generates motor representations and maintains readiness for movement (Cunnington *et al*. 2005). In addition to these earlier works, our current results showed that a prolonged motor planning process in the human non-primary motor cortex impaired subsequent motor command and execution: the more sustained the preparatory signals in the PM and SMA, the slower the RTs.

More importantly, our results further showed that in patients with electrode implantations in both PM and M1, the duration of PM HGB activation positively correlated with the peak latency of M1 HGB power: the longer the motor planning lasted in PM, the later the motor command took place in M1 (Fig. 5D). To investigate the relationship between the firing rate of the motor cortices and the RTs, most of the previous animal research dissociated motor preparation and execution by adopting the delayed reaching paradigm (Churchland, Santhanam*, et al.* 2006; Churchland, Yu*, et al.* 2006; Rickert et al. 2009; Afshar *et al*. 2011; Ames *et al*. 2014; Hasegawa *et al*. 2017). For example, in the delayed reaching task, the firing rate of the PM region during motor preparation can predict the following movements in monkeys (Churchland, Santhanam*, et al.* 2006; Churchland, Yu*, et al.* 2006; Rickert *et al*. 2009; Afshar *et al*. 2011; Ames *et al*. 2014). Critically, the RTs, rather than the movement type, can be predicted by the firing rate pattern in the PM region during motor preparation (Afshar *et al*. 2011; Kaufman *et al*. 2016). When the PM firing rate variability is higher during motor preparation, the RTs of the movement will be longer (Churchland, Santhanam*, et al.* 2006; Churchland, Yu*, et al.* 2006). However, it remains unclear whether these results can be generalized to humans. Previous evidence from human and monkey intracranial EEG research has shown a close relationship between the neuronal firing rate and the high-gamma oscillation of local field potential (LFP). Specifically, the neuronal firing rate is strongly correlated with HGB power, temporally and, more importantly, on a trial-by-trial basis, in the sensorimotor cortex (Ray, Crone, et al. 2008; Ray, Hsiao, et al. 2008; Ray and Maunsell 2011) and the visual cortex (Rash et al. 2008; Keeley et al. 2019). Together with previous evidence, the neuronal firing rate (Churchland, Santhanam*, et al.* 2006; Churchland, Yu*, et al.* 2006; Churchland and Shenoy 2007) and the duration of HGB power activation in the PM during motor preparation observed in the present study represent neural signatures of motor preparation in the higher-order motor cortex. The higher firing rate and the more sustained HGB power in the PM region lower the efficiency of motor planning, which subsequently delays motor command in terms of prolonged peak latency of HGB power in the M1 region and accordingly results in slower RTs.

Direct comparisons between the peak latency of HGB power in the three regions showed that the PM and SMA peaked approximately 300 ms after the cue (Fig. 5A and sFig. 6A upper and middle panel), confirming an earlier motor planning function. In contrast, M1 peaked approximately 400 to 800 ms after the cue, and the M1 HGB power peak was strictly time-locked to movement onset (Fig. 5A and sFig. 6A lower panel), confirming a later motor command function. The present results are consistent with previous evidence showing sequential HGB activation of PM and M1 during the preparation of finger movements (Sun et al. 2015). The sequential activation from PM to M1 implied a possible pathway via which information was transmitted from the non-primary motor cortices to the primary motor cortex. The present pairwise phase consistency results showed that the pairwise HGB coherence between M1, PM, and SMA peaked approximately 300 ms after cue onset, i.e., during the preparatory phase (Fig. 6). Moreover, for the three pairwise connections and the contralateral and ipsilateral finger movements, the more sustained the HGB coherence, the slower the finger movements (Fig. 6), which is highly consistent with the HGB power results in the PM and SMA (Fig. 4A, B, D, and E lower panel). Therefore, motor planning is associated with the local HGB power within the higher-order motor cortices in the PM and SMA and interregional connections between the non-primary and primary motor cortices.

Some limitations are worth considering. The limitations of iEEG recording led to sparse and spatially biased sampling. Furthermore, the placement of electrodes is determined clinically and, once implanted during surgery, cannot be altered—unlike in animal studies, where researchers may penetrate the cortex multiple times until responsive neurons are identified. In the current study, only four patients had responsive electrodes simultaneously in PM, SMA, and M1.

The structural connectivity between M1, PM, and SMA was traced in the monkeys’ motor system (Ninomiya et al. 2019). The non-primary motor cortex is connected directly to the primary motor cortex in both hemispheres (Luppino et al. 1993; Liu *et al*. 2002). Functionally, the interplay between the non-primary and primary motor cortex is vital for motor preparation (Espenhahn et al. 2017). For example, during motor preparation, PM-M1 excitatory interactions facilitate voluntary movements by increasing corticospinal excitability (Cote et al. 2017; Neige et al. 2021), and patients with motor dysfunction show malfunctions in the connection between the M1, PM, and SMA (Piramide et al. 2022). For the first time, the present results delineated a relatively complete picture of how the primary and the non-primary motor cortices work together to achieve efficient finger movements (Fig 7): Motor planning in the higher-order motor cortices needs to end fast, both in terms of shorter local HGB activations and interregional communication/connections, so that movement execution in M1 can be efficiently initiated, i.e., RTs will be faster.

**Fig. 7.**
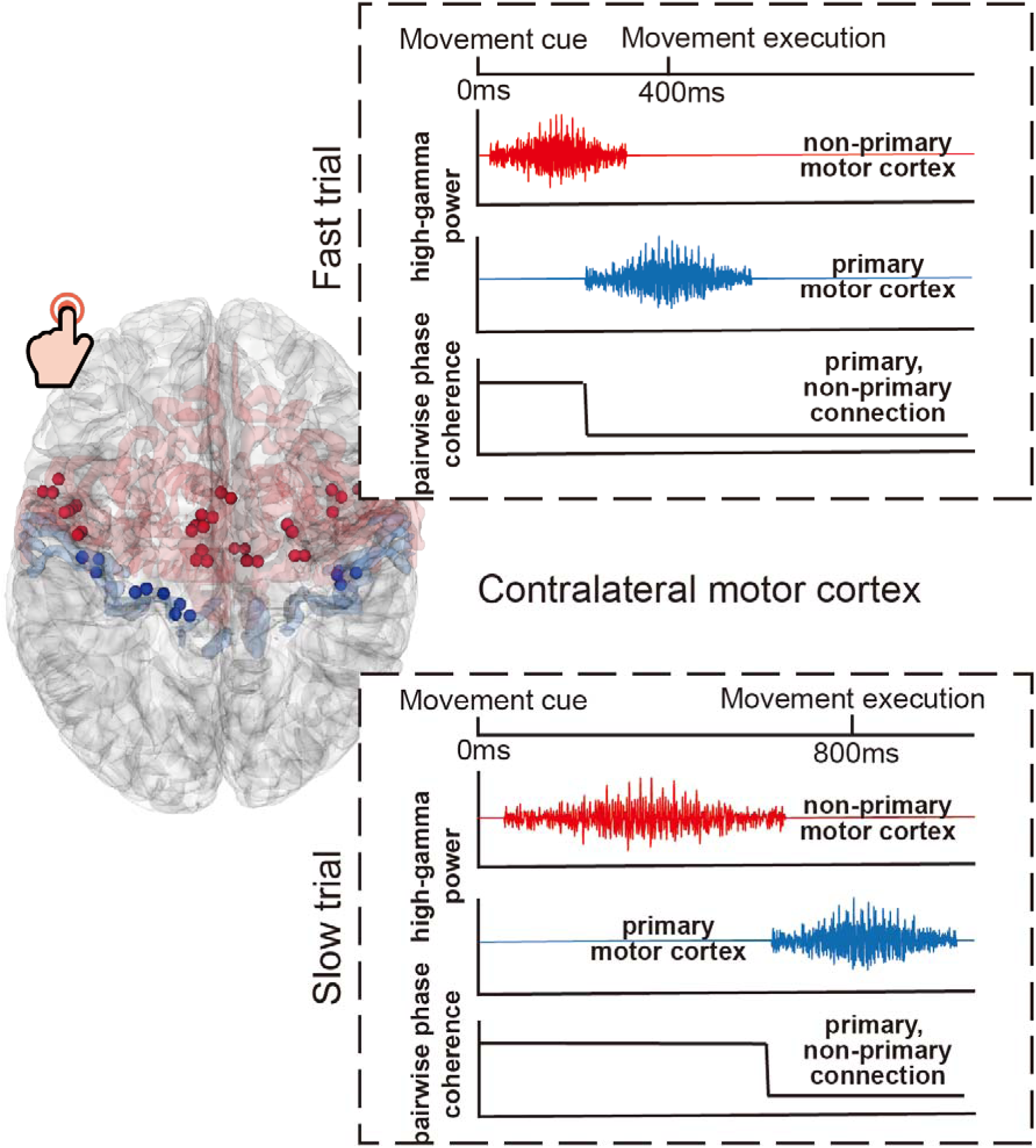
Schematic map of how the HGB power and HGB PPC in the primary and non-primary motor cortex affect reaction times during contralateral finger movement. Both motor planning activity in the non-primary motor cortex (red signals) and non–primary–primary motor coherence (black signals) end earlier in the fast trials than in the slow trials so that motor execution (blue signals) initiates earlier in the fast trials than in the slow trials. In the 3D glass brain on the left, the primary motor electrodes are marked in blue, while the non-primary motor electrodes are marked in red.

## Supporting information

sFig

## Acknowledgements

This work was supported by a grant to SD and GRF by the Deutsche Forschungsgemeinschaft (DFG, German Research Foundation) - Project-ID 431549029, SFB 1451. SD acknowledges further funding by the DFG (491111487). QC is supported by grants from the Natural Science Foundation of China (31871138, 32071052) and the MOE Project of Key Research Institute of Humanities and Social Sciences in Universities (22JJD190006). JX is supported by grants from the Guang Dong Basic and Applied Basic Research Foundation (No. 2021A1515110125) and by the Grant from Shenzhen Science and Technology Innovation Committee (No. JCYJ20190806171209030).

