## Supplementary material for "Earlier finish of motor planning in the premotor cortex predicts faster motor command in the primary motor cortex: human intracranial EEG evidence": sFig

### Supplementary materials:

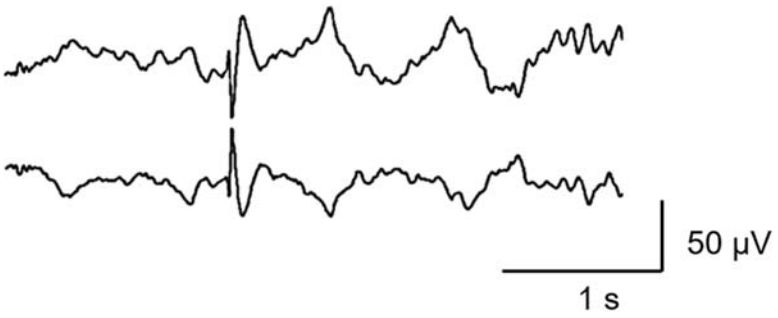


**sFig. 1 Example of interictal epileptiform discharge (IED).** If any trial shows the IED, the whole trial will be deleted from further analyses.

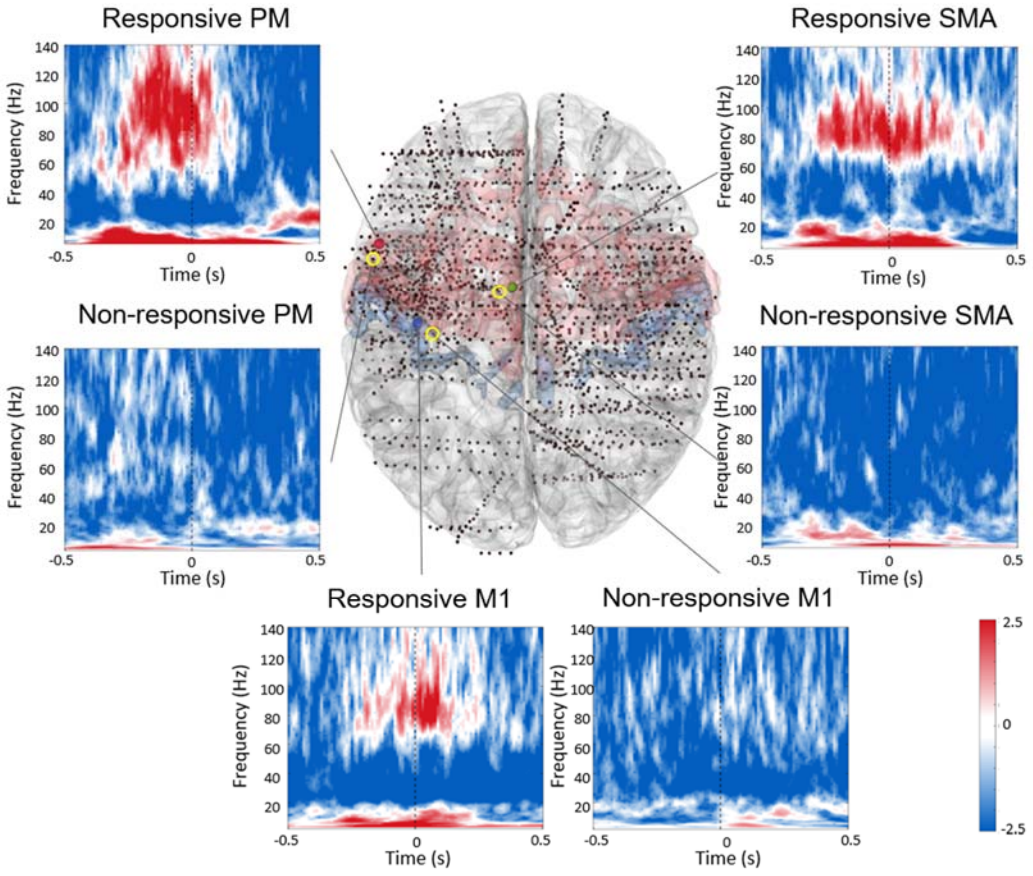


**sFig. 2 Examples of responsive and nonresponsive PM, SMA and M1 electrodes in bilateral finger movement trials.** Group-level time-frequency results in example electrodes of the PM (red electrode), SMA (green electrode), and M1 (blue electrode) during the bilateral finger movement condition are displayed. Time zero (marked by the black dashed line) indicates the initiation of finger movement. The electrodes within the yellow circles showed neither increased gamma nor decreased

beta power around the finger movement. These electrodes were defined as nonresponsive electrodes and were excluded from further analyses.

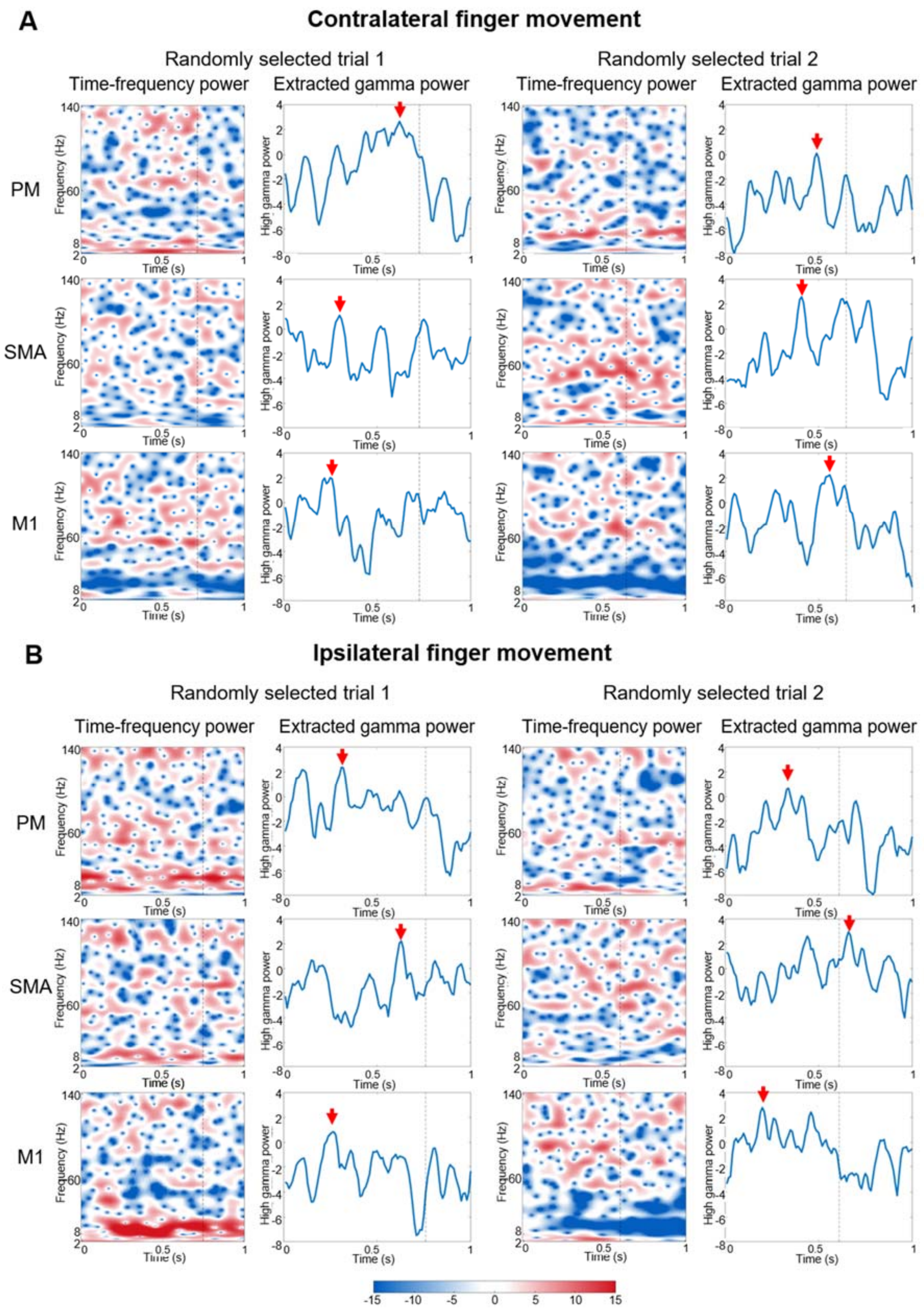


**sFig. 3 Time-frequency and extracted gamma power in the PM, SMA and M1 in randomly**

**selected single trials during the contralateral (A) and ipsilateral (B) finger movement.** In both A) and B), the time-frequency power spectrum and the high-gamma (60-140 Hz) power were extracted in two randomly selected single trials. The black dashed lines mark the onset of finger movements. The red arrows indicate peak activation of the high-gamma power, which correspondingly defines peak latency and area under the curve. It is evident that nonspecific noise can easily contaminate the determination of high-gamma power peaks in single trials and can be easily contaminated by nonspecific noise.

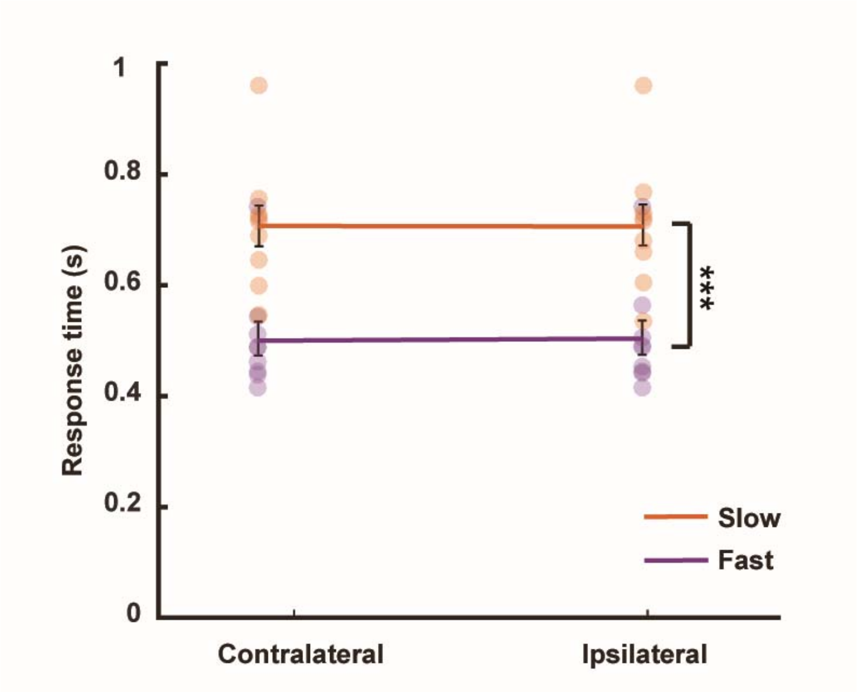


**sFig. 4 Behavioral results.** The mean reaction times (RTs) are displayed as a function of the side of the response hand (contralateral vs. ipsilateral) and the response speed (fast vs. slow). A 2 x 2 repeated-measures ANOVA showed that the main effect of response speed was the only significant effect, $F_{(1, 8)} = 286.243$, $p < 0.01$, indicating a significant RT difference between the fast and slow trials for both contralateral and ipsilateral responses. The dots represent the average reaction time of each patient. Error bars indicate the standard error of the mean.

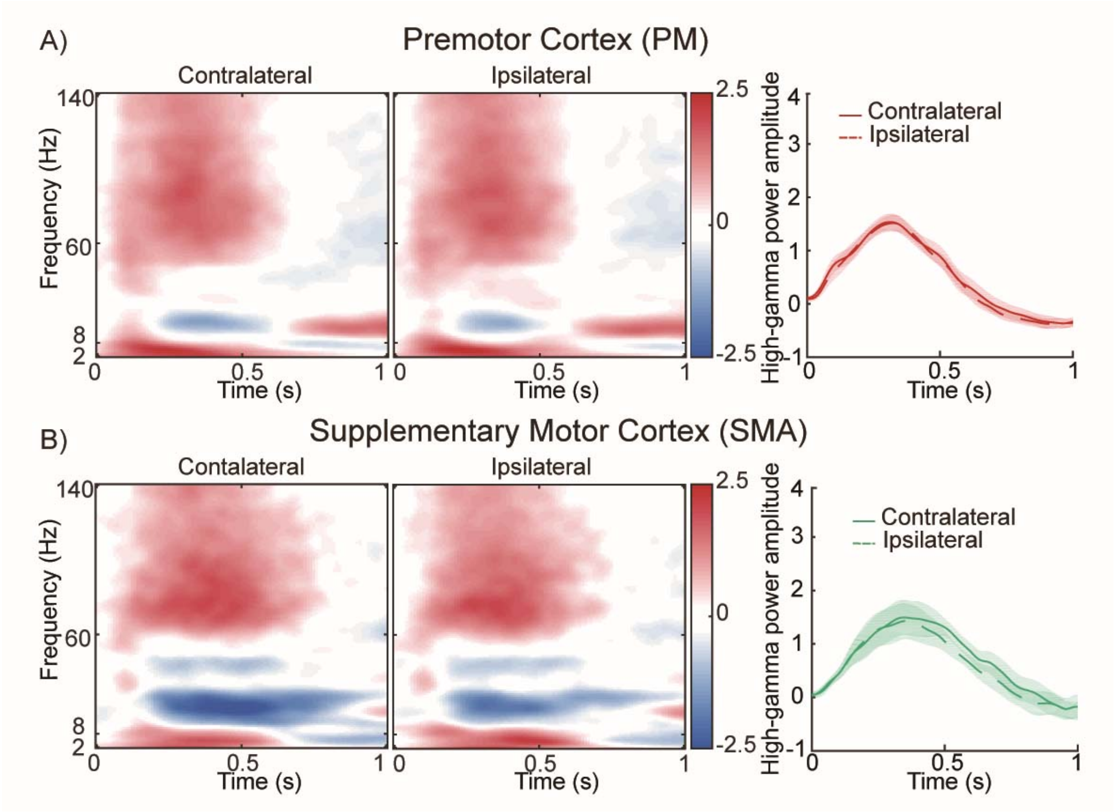


**sFig. 5 Time-frequency results of PM and SMA. A)** Group-level time-frequency results of the premotor cortex. **B)** Group-level time-frequency results of the supplementary motor cortex. In **A)** and **B)**, the temporal profiles of high-gamma power (60-140 Hz) are shown as a function of the contralateral movement (red/green solid lines) and ipsilateral movement (red/green dashed lines) trials. Time ticket zero marks the onset of the stimulus. The shadow around each solid/dashed line indicates the standard error of the mean.

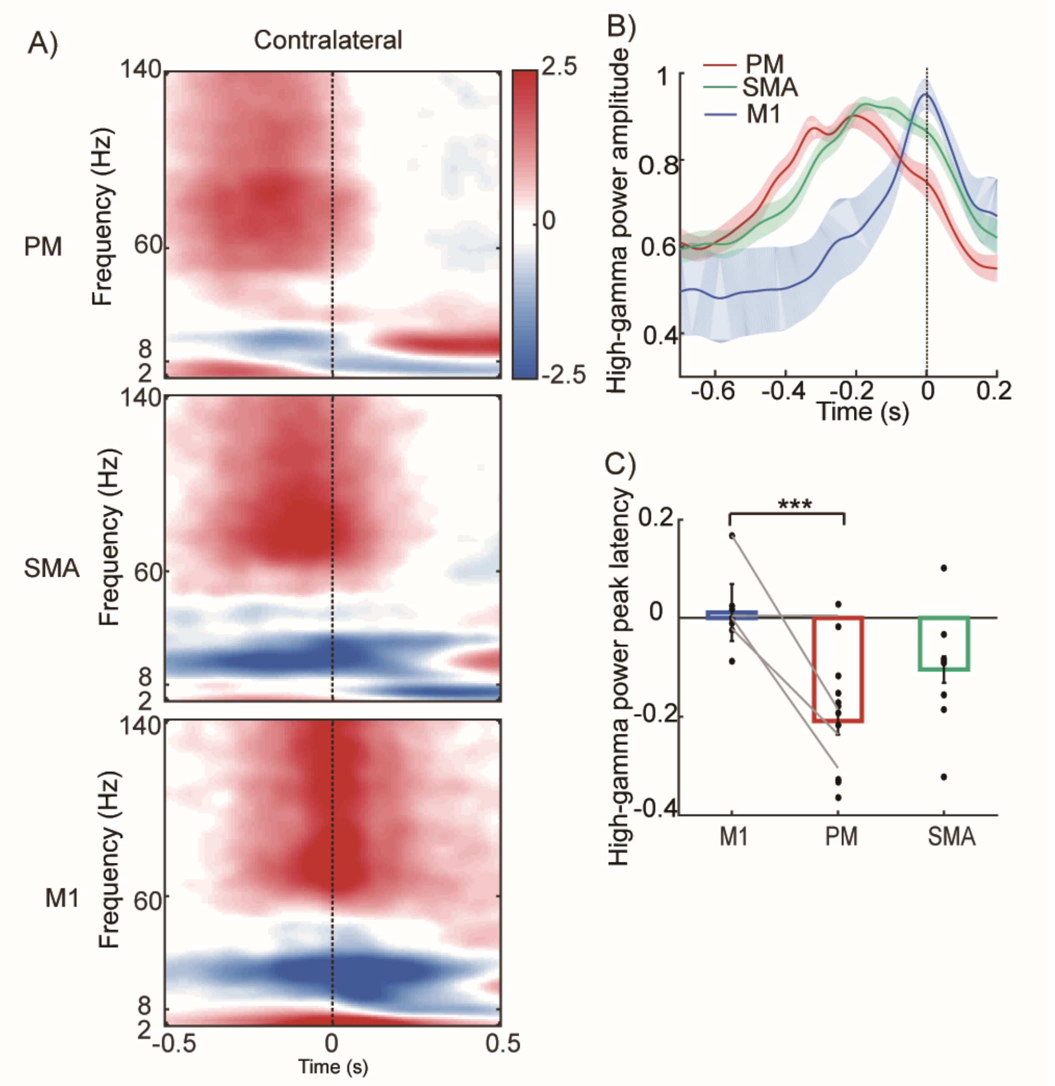


**sFig. 6 Sequential activations of PM, SMA, and M1 when time-locked to the response. A)** For the contralateral movement conditions, the power of the high-gamma band was activated sequentially. **B)** The high-gamma power was extracted and averaged for PM, SMA, and M1. The shadow around each line indicates the standard error of the mean (SEM). **C)** The time to power peak of the high-gamma band was submitted to a one-way ANOVA of each electrode ($F_{(2, 30)} = 8.392$, $p < 0.01$, post hoc *t* tests revealed a significantly shorter HGB peak latency in PM than M1, $t_{(20)} = 4.299$, $p < 0.01$). Each gray line connecting two data points represents the average peak latency of one patient. These results were based on four patients with electrodes implanted simultaneously in the PM, SMA, and M1. *** $p < 0.01$; with Bonferroni correction. The shadow around the line in B and the error bars in C indicate SEM. The black dashed line represents the movement onset of the hand.
